# The behavioral phenotype of early life adversity: a 3-level meta-analysis of preclinical studies

**DOI:** 10.1101/521245

**Authors:** V Bonapersona, J Kentrop, CJ Van Lissa, R van der Veen, M Joëls, RA Sarabdjitsingh

## Abstract

1

**Background:** Altered cognitive performance has been suggested as an intermediate phenotype mediating the effects of early life adversity (ELA) on later-life development of mental disorders, e.g. depression. Whereas most human studies are limited to correlational conclusions, rodent studies can prospectively investigate how ELA alters cognitive performance in a number of domains. Despite the vast volume of reports, no consensus has yet been reached on the *i)* behavioral domains being affected by ELA and *ii)* the extent of these effects.

**Methods:** To test how ELA (here: aberrant maternal care) affects specific behavioral domains, we used a 3-level mixed-effect meta-analysis, a flexible model that accounts for the dependency of observations. We thoroughly explored heterogeneity with MetaForest, a machine-learning data-driven analysis never applied before in preclinical literature. We validated the robustness of our findings with substantial sensitivity analyses and bias assessments.

**Results:** Our results, based on >400 independent experiments, yielded >700 comparisons, involving ~8600 animals. Especially in males, ELA promotes memory formation during stressful learning but impairs non-stressful learning. Furthermore, ELA increases anxiety and decreases social behavior. The ELA phenotype was strongest when *i)* combined with other negative experiences (“hits”); *ii)* in rats; *iii)* in ELA models of ~10days duration.

**Conclusion:** Prospective and well-controlled animal studies demonstrate that ELA durably and differentially impacts distinct behavioral domains. All data is now easily accessible with MaBapp (https://osf.io/ra947/), which allows researchers to run tailor-made meta-analyses on the topic, thereby revealing the optimal choice of experimental protocols and study power.

## 2 Introduction

Early life adversity (ELA) is a consistent risk factor of psychiatric disorders (1, 2), and it is regularly associated with poorer cognitive outcomes later in life (3–5). Indeed, impaired cognitive processing is a prominent feature of psychopathologies (3, 6, 7), e.g. dysregulated contextual memory in post-traumatic stress disorder (8) or social cognition in schizophrenia (9). ELA may therefore alter cognitive development, thereby resulting in behavioral abnormalities that may render individuals more vulnerable to psychiatric disorders (10).

To investigate exactly how ELA affects cognitive processing, rodent models are a valuable resource: they complement human studies by in-depth and thorough investigations of otherwise hard-to-study mechanisms. In animal experiments, genetic and environmental influences can be more precisely controlled and experimentally varied than in humans (11). Furthermore, prospective designs are more feasible. For example, rodent studies have disentangled the different components of mother-pup interaction, a critical factor of early development across mammalian species (12, 13). This has helped uncover links between disturbed maternal care and disturbed emotional and cognitive functioning later in life, implicating the stress system (14) and “hidden regulators” (15).

Rodent studies have also highlighted paradoxical ELA effects on cognitive abilities. For instance, Benetti *et al*. (16) reported that rats with a history of ELA had impaired memory in the object recognition task. Conversely, Champagne *et al*. (17) reported that ELA mice display increased memory in a fear conditioning paradigm. Both tests have historically been used as memory tasks, albeit in a non-stressful and stressful context respectively. Possibly, the equivocal results are due to different underlying biological mechanisms (e.g. learning in stressful vs non-stressful situations) or pertain to the divergent methodology used (e.g. type of test or ELA model, species, experimenters, labs). A few studies have investigated the latter by testing the same animals in different memory tasks (18–21). Although these studies favor the former explanation, the limited amount of animals used (22)–alongside the heterogeneous methodology– prohibits firm conclusions.

To address this conundrum, we here carried out a large-scale 3-level meta-analysis of all peer-reviewed preclinical literature on the subject, and tested the hypothesis that ELA (particularly aberrant maternal care) differentially affects specific behavioral domains in adulthood. We focused on memory formation after stressful or non-stressful learning, anxiety and social behavior, given their relevance for psychopathologies. We addressed (potential) sex-differences by investigating males and females separately. Furthermore, we tested whether the presence of multiple hits (23) (e.g. other negative life experiences) amplified ELA effects. Finally, we applied a novel, machine-learning based analysis (MetaForest(24)) to identify the most important moderators of ELA effects on behavior.

Based on this comprehensive analysis, we evaluate the translational potential of ELA rodent models. With the aid of a specially developed web-based tool MaBapp *(Meta-Analysis of Behavior application)* (https://osf.io/ra947/), interested researchers can perform their own meta-analysis and retrieve valuable *ad hoc* information for experimental design and power calculations.

## 3 Methods

We adhered to SYRCLE’s guidelines (25)(26), and to the PRISMA (27) reporting checklist. To ease reading of the methodology, definitions of technical terms are provided in Supplemental Methods (S1.1). A summary of the general approach can be found in Figure 1.

**Figure 1.**
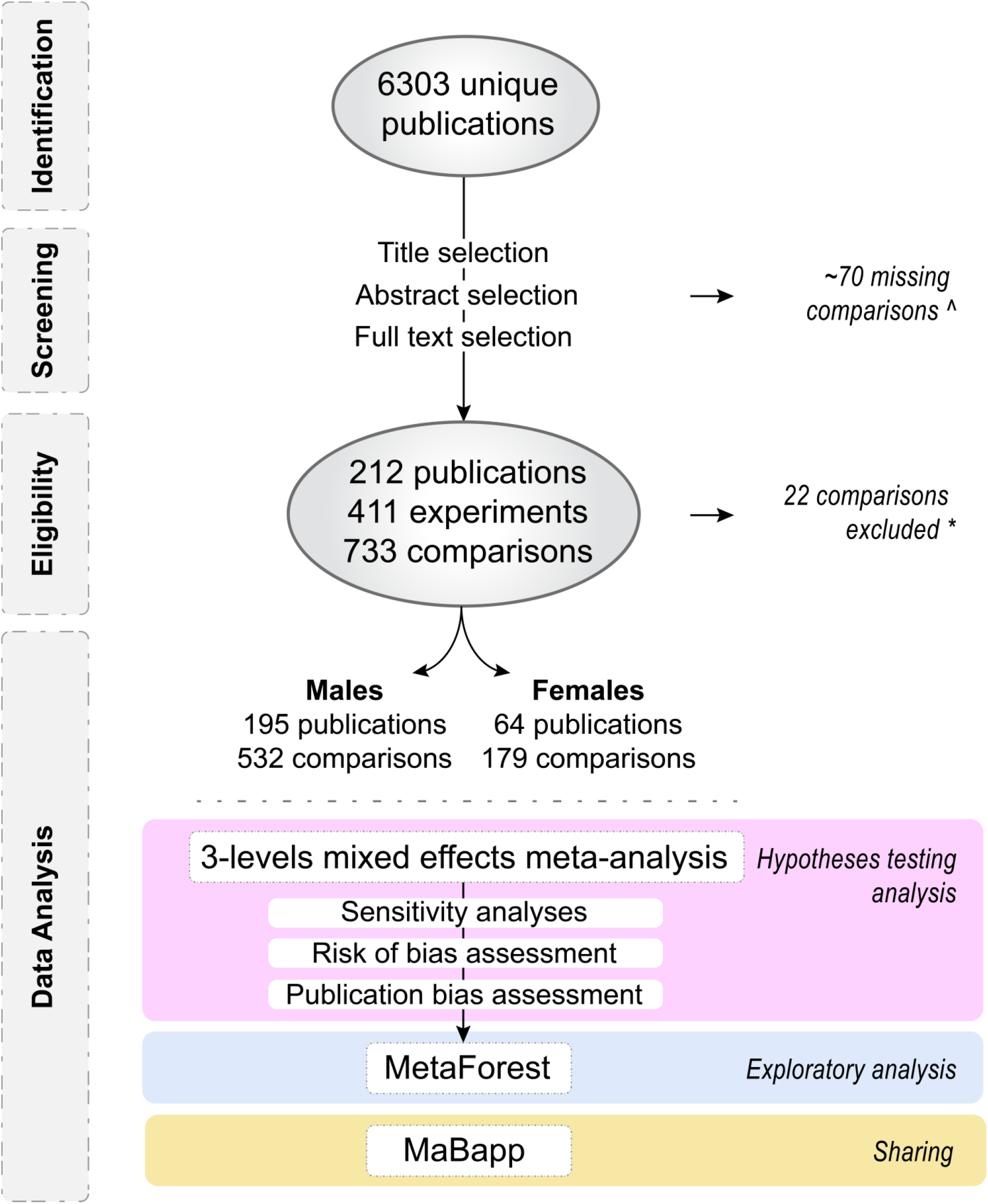
Flow chart of study selection and analysis. ^ = estimation of missing comparisons (S2.1); * = comparisons excluded from the meta-analysis due to controversial behavioral domain categorization (S2.2)

### 3.1 Search Strategy

The electronic databases PubMed and Web of Science (Medline) were used to conduct a comprehensive literature search on *the effects of ELA on behavior* on December 6^th^ 2017. The search string was constructed with the terms “behavioral tests”, “ELA” and “rodents” (S1.2).

Prior to the beginning of the study, four experts were consulted. After elaborate discussions they agreed upon *i)* the selection of tests and related outcomes (S1.3), *ii)* their classification into behavioral domains (S1.3) and *iii)* the definition of multiple hits (S1.4). The results of each individual test, independent of categorization, are available for consultation on MaBapp (Section 5.1). Studies’ titles and abstracts were screened independently by two researchers (VB & JK) and selected if the inclusion criteria were met (S1.5). Studies’ inclusion was performed blinded to the studies’ results. In case of doubt, the full text was inspected. Any disagreement was resolved by greater scrutiny and discussion.

To limit subjectivity in the data gathering and entry process, data from eligible studies were extracted in a standardized dataset alongside its explanatory codebook (https://osf.io/ra947/).

For each individual comparison, we calculated *Hedge’s* G (28), a standardized mean difference with a correction for small samples (29). S1.6 details the extraction of statistical information as well as handling of missing values. We estimated the summary statistics of data presented only graphically with *Ruler for Windows* (https://a-ruler-for-windows.en.softonic.com/), of which we previously validated the accuracy (30). If the data was not reported in any format (or other crucial information was missing e.g. sex), we contacted two authors per manuscript published after 2008 (response rate 52.6%). If no answer was received within two months and after a reminder, the authors were considered not reachable, and the comparison was excluded.

### 3.2 Meta-analysis: research questions and statistical approach

To avoid possible biases, the experimenter (VB) was blinded to the ELA effects while coding the analysis. This was achieved by randomly multiplying half of the effect sizes by -1.

#### 3.2.1 Hypothesis-testing

We built a 3-level mixed effect meta-analysis with restricted maximum likelihood estimation. In our experimental design, the 3 levels correspond to variance of effect size between 1) animals, 2) outcomes and 3) experiments. This approach accounts for the violation of the assumption of independency when the data is collected from the same animals (30–32), thereby improving the robustness of the conclusions drawn. We included “domains” and “hits” as moderators in order to address the following two research question: 1) *what are the effects of ELA on each behavioral domain?*; 2) *are the effects enhanced if the animals experienced multiple hits?*. Since both questions were answered with the same model, effect sizes were estimated only once.

We ensured that all behavioral measurements were in the same theoretical direction by multiplying – whenever necessary – the effect sizes by -1 (29)(S1.3). Although this was essential for the model estimation, we here report effect sizes in a more interpretable manner: an increase in Hedge’s G signifies an enhancement of the behavioral domain under study (e.g. more anxiety, better memory).

We conducted several sensitivity analyses (S1.7) to assess the robustness and consistency of our conclusions. We examined whether the quality of the studies affected the estimation of the results by dissecting the influence of reporting bias, blinding, randomization and study power. Furthermore, we thoroughly investigated influential and outlying cases (33) according to multiple definitions (S1.7).

To compensate for methodological limitations, we tested the presence of publication bias with various qualitative/quantitative methods (S1.8), and quantified its influence with fail-safe N (34) and trim-and-fill analyses (35)(S1.8).

Risk of bias was evaluated with SYRCLE’s assessment tool (36), where we distinguished between study-level and outcome-level biases (27). Lack of reporting of experimental details was scored as an *unclear* risk of bias.

Heterogeneity was assessed with Cochrane Q-test (32) and I^2^, which was estimated at each of the 3-levels of the model to determine how much variance could be attributed to differences within (level 2) or between experiments (level 3) (37). Estimates of explained variance can be positively biased when based on the data used to estimate the model (38). For this reason, we used 10-fold cross-validation to obtain an estimate of how much variance our model might explain in new data. This cross-validated estimate of R^2^ (R_cv_^2^) is robust to overfitting and provides evidence for the results’ generalizability. P-values were corrected with Bonferroni for family-wise error rate (each research question considered as a separate family of tests) to limit capitalization on chance. Since we expected the amplitude of effect sizes to differ between sexes (39, 40), we considered males and females as two separate datasets.

#### 3.2.2 Exploratory analysis

We used MetaForest (24), an novel exploratory approach to identify the most important moderators of the ELA effects on behavioral domains. This innovative, data-driven technique adapts random forests (a machine learning algorithm) for meta-analysis, by means of bootstrap sampling. MetaForest ranks moderators based on their influence on the effect size.

Preclinical experiments often adopt diverse protocols. Although this can be an advantage (41), in a meta-analysis it induces substantial heterogeneity. Therefore, we classified the published experimental protocols in >30 standardized variables with the intent to identify potential methodological sources of heterogeneity. Based on theoretical importance, we selected 26 of these moderators for inclusion in the MetaForest analysis. We used 10-fold cross-validation (S1.9) to determine the optimal tuning parameters that minimized RMSE: uniform weighting, 4 candidate moderators at each split, and a minimum node size of 2. The marginal bivariate relationship of each moderator with effect size was averaged over the values of all other moderators (S1.9). Residual heterogeneity was estimated with tau^2^ (S1.9).

Lastly, we created MaBapp (https://osf.io/ra947/) for anyone to perform their own meta-analysis on the topic by selecting their favorite characteristics (Section 5.1).

Analyses were conducted in R (version 3.5.1) (42), using the following packages: 1) *metafor* (28) for conducting the analysis, 2) *metaforest* (43) for data exploration, 3) *shiny* (44) to create MaBapp, and 4) *dplyr* (45) for general data handling. For further specifications about the analysis, the R script is made available (https://osf.io/ra947/).

## 4 Results

### 4.1 Studies selection and characteristics

In total ~8600 animals (age_weeks_ median[IQR]=12[4]; proportion rats=68%) were included in the analysis, 77.7% of which were males. Anxiety was the behavioral domain most investigated (48.8%), elevated plus maze the most popular test (14.3%), and maternal separation the ELA paradigm most often used (48.9%). For additional descriptive information on study characteristics, see S2.3.

Although no publication reported on all SYRCLE’s potential bias items, 41 publications (19.3%) were blinded as well as randomized, and overall we estimated a risk of bias of 3[1] (median[IQR]) on a 10 points scale (S2.4). Lastly, at a systematic review level (S2.5), 68.5% of comparisons were either not-significant (n_comp_=386) or the result could not be directly interpreted from the information provided (n_comp_=117).

### 4.2 ELA effects are pronounced in males and with “multiple hits”

The effect sizes included ranged between −6.4 and 6.1 (mean[SD]=.29[1.06]), with 95% of comparisons between −2 and 2. Sample size ranged between 6 and 59 animals (mean[SD]=22[7.8]), and differed <20% between control and ELA groups in 90% of the cases (estimation).

When qualitatively comparing sexes, the effects of ELA were more evident in males than in females. **Male** rodents with a history of ELA displayed increased anxiety (HedgesG[95%CI]=.278[.165,.39], z=4.819, p<.000), improved memory after stressful learning (HedgesG[95%CI]=.283[.141,.425], z=3.9, p<.000), impaired memory after non-stressful learning (HedgesG[95%CI]=-.594[-.792,-.395], z=-5.86, p<.000) and decreased social behavior (HedgesG[95%CI]=-.614[-.88,-.348], z=-4.521, p<.000, Figure **Error! Reference source not found**.A, S2.6). We were unable to confirm any effect of ELA on behavior in females, although directionality was generally comparable in both sexes (**Error! Reference source not found**.B, S2.7).

Overall, the presence of multiple hits intensified the effects of ELA in **males** (HedgesG[95%CI]=.222[.018,.426], z=2.131, p=.033) yet marginally in **females** (HedgesG[95%CI]=.297[-.003,.596], z=1.939, p=.052). Although these enhancing effects were not significant at a single-domain level (posthoc analysis) (Figure 2C-D, S2.6/S2.7), memory after non-stressful learning was the most impacted domain in **males** (difference in Hedge’s G=.435, z=2.156, p=.124) as well as in **females** (difference in Hedge’s G=.565, z=2.234, p=.102).

**Figure 2.**
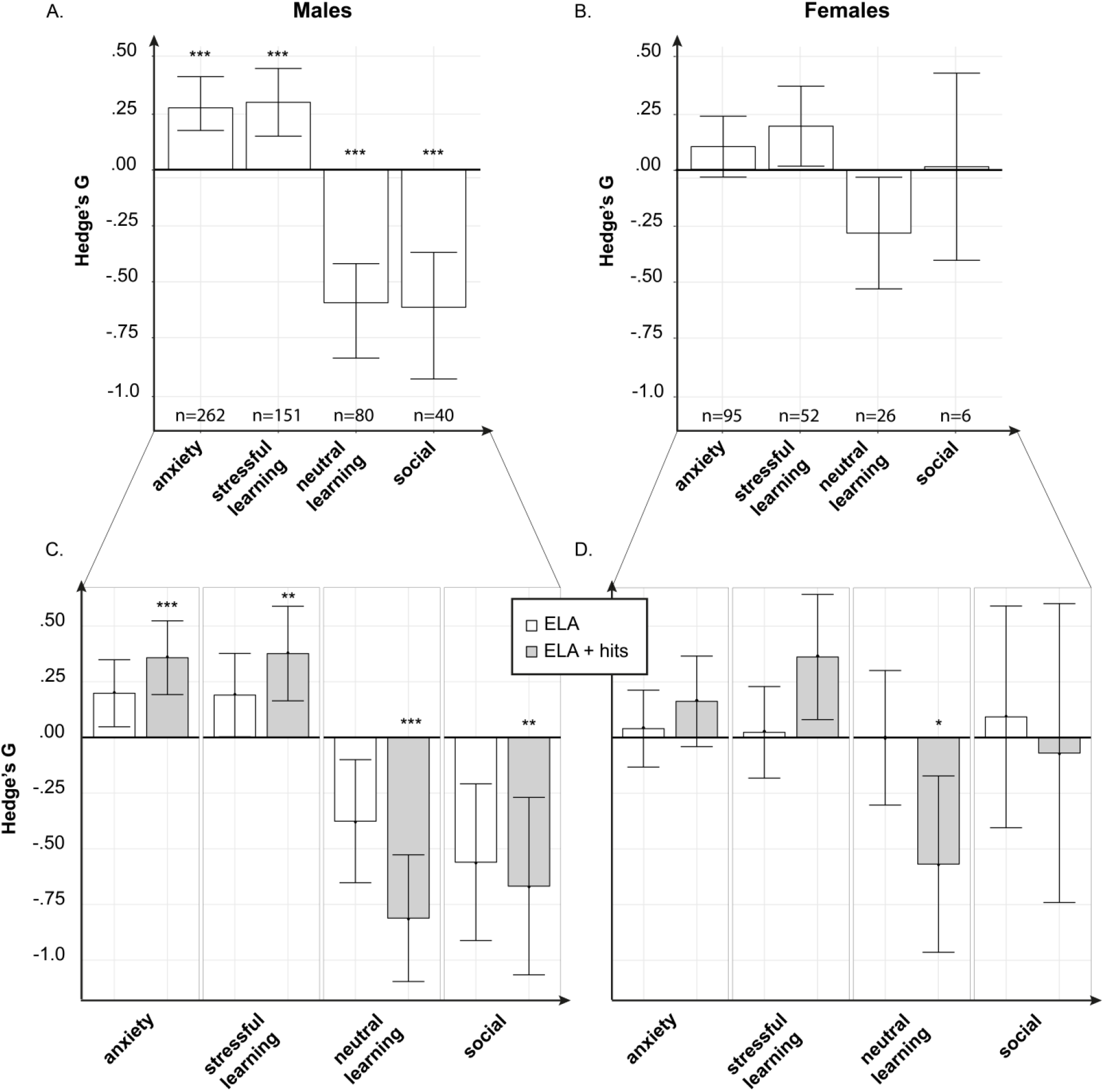
Effects of ELA on behavioral domains in males (**A**) and females (**B**), and the role of multiple hits (in addition to ELA, grey bars) compared to only ELA (white bar) in mediating these effects (males: **C**, females: **D**). Each bar represents the size of the effect (*Hedge’s G*, standardized mean difference) of the ELA manipulation when comparing a control and an experimental group. * = p<.05, ** = p<.01, *** = p<.001

#### 4.2.1 Sensitivity analyses and publication bias

Qualitative evaluation of funnel plot asymmetry suggested the presence of publication bias, which was confirmed by Egger’s regression and Begg’s test (S2.8). Nonetheless, fail-safe N as well as trim-and-fill analyses confirmed that – albeit present – publication bias is unlikely to distort the interpretation of the results (S2.8). Furthermore, the robustness of the male and female models was confirmed by several sensitivity analyses (S2.9).

### 4.3 Exploration of Moderators

Although the models of the hypotheses-testing analysis described a significant proportion of variance (R_cv_^2^_males_=.026, R_cv_^2^_females_=.03), substantial heterogeneity was recorded in both models (**males**: Q(524)=1763.118, p<.000; **females**: Q(171)=326.93, p<.000, S2.10). This was not surprising due to the diversity of publications included in the meta-analysis.

To investigate the source of the heterogeneity, we used MetaForest, a new statistical technique that ranks moderators (Figure 3A) based on their predictive value. These can roughly be divided in 4 groups, describing: *i)* characteristics of the animals (e.g. origin of the breeding animals (Figure 3B) and species investigated (Figure 3C)), *ii)* ELA model used (e.g. type of model and duration of ELA (Figure 3D-E)), *iii)* outcome measures (e.g. domain and test used), and iv) potential bias (e.g. blinding and randomization). MetaForest confirmed that the selected moderators account for a substantial portion of the variance (R_cv_^2^[SD]=.12[.09]).

**Figures 3.**
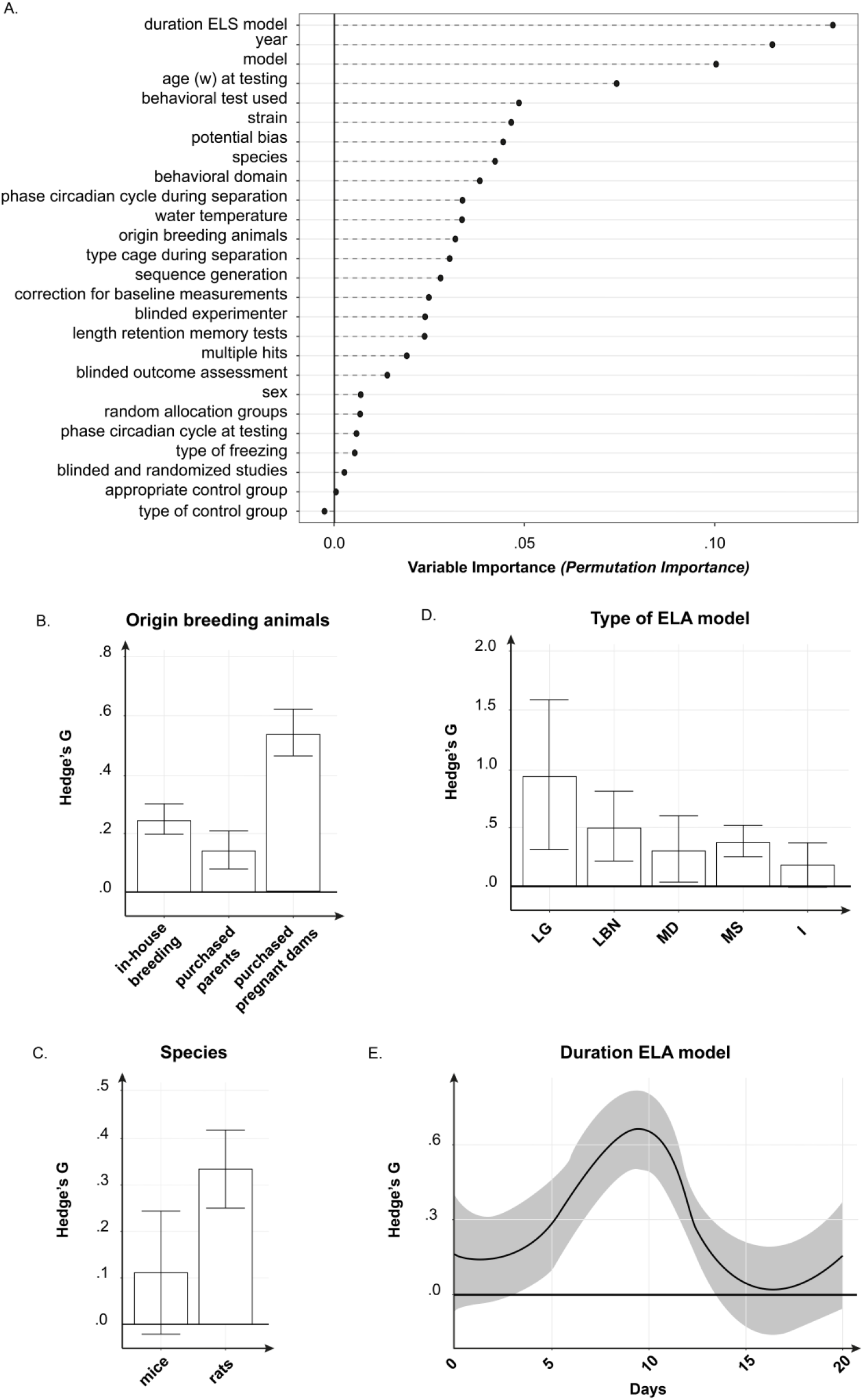
Exploratory MetaForest analysis. (**A**) Rank moderators’ importance. Variable/permutation importance is a measure of how strongly each moderator explains differences in effect size, capturing (non-)linear relationships as well as higher order interactions. For information about MetaForest’s partial dependence plots, see S2.11. Effect sizes distinguished by origin of the breeding animals (**B**), species (**C**), type of ELA model (**D**) and duration of ELA (**E**). Results are expressed as Hedge’s G[95%CI]. The usefulness of this exploration can be best appreciated with the aid of MaBapp. For example, the overall estimate of the effects of ELA on anxiety is Hedge’s G=.24. However, if we select only the LBN model, the effect size rises to .37. If we combine LBN and rats, the effect size further rises to .60. If we then select only elevated plus maze as respectively behavioral test, the effect size rises to .81. LG = licking-and-grooming, LBN = limited bedding and nesting, MD = maternal deprivation, MS = maternal separation, I = isolation.

Offspring of dams purchased pregnant had larger effect sizes than offspring bred in the own facility (Figure 3B). Rats had overall larger effect sizes than mice (Figure 3C). Concerning ELA models (Figure 3D), selecting the extremes of natural variation (licking-and-grooming model) yielded the strongest phenotype. Lastly, effect sizes appeared to be maximal with a 10 days’ ELA duration (Figure 3E).

## 5 DISCUSSION

In this study, we substantiate that adversities early in life profoundly and lastingly change rodent behavior. Due to low power (22) and heterogeneous methodologies, results at a single-study level are often inconclusive and difficult to interpret. Here, by adopting a meta-analytic approach, we provide extensive evidence that ELA has differential effects on memory: it enhances memory if learning occurs in a stressful situation, but it hampers learning under non-stressful circumstances. Furthermore, ELA increases anxiety and decreases social behavior, particularly in males. In line with the multiple-hits hypotheses (23, 46), the effects are amplified if the animals experience other stressful life events (e.g. prenatal stress due to transport of pregnant females). These results are independent of the type of ELA or behavioral test used, and are remarkably similar to what was reported at a correlational level in humans (47, 48). Altogether, our results point to a clear and robust phenotype of ELA in four behavioral domains and complement the human literature by supporting a causative role of ELA in altering behavior, which may predispose individuals to precipitate symptoms of psychiatric disorders.

### 5.1 Methodological considerations

The lack of sufficient power to detect experimental effects is an emerging issue in preclinical literature (22, 30) that seriously hampers research interpretation (22). As a consequence, results from single-studies are useful for hypotheses generation but do require replication. Indeed, in our study the majority of comparisons (68.5%) was not-significant at a systematic review level, but the effects were significant when analyzed meta-analytically. Although still sparsely applied to preclinical studies, statistical tools such as meta-analyses can therefore be very useful to substantiate conclusions from animal data and translate them more reliably to patients (49).

In this project, we intertwine this concept with state-of the-art statistical methodology, adopting an approach never used in preclinical studies. Firstly, our meta-analysis was built with a 3-level model (32), which allows for a more robust estimation of the effects by accounting for the dependency of same-animal’s data (30, 50). Secondly, a leading strength of preclinical meta-analyses is the systematic exploration of heterogeneity (49). Instead of the standard subgroup/meta-regression approach, we opted for an exploratory analysis using MetaForest (24), a newly developed technique that ranks moderators’ importance by adapting the machine learning algorithm *random forests* to summary-statistics’ data. A major strength of MetaForest is its robustness to overfitting, and its ability to accommodate non-linear effects, as shown by the impact of ELA duration on effect sizes (24).

Thirdly, we extensively coded potential (biological and experimental) moderators. Although possibly relevant moderators were not included due to insufficient reporting (e.g. temperature during separation (51), cross-fostering (52), culling (53)), this dataset treasures relevant information for future experimental designs. To facilitate others to exploit this dataset, we created **MaBapp** (https://osf.io/ra947/), a web-based app with a user-friendly interface through which anyone can perform his/her own meta-analysis on the topic of ELA and behavioral domains. Within the app, a wide variety of features can be selected, such as ELA models and their components (e.g. type, timing, predictability), behavioral tests used, age and sex of the animals, etc. Based on the characteristics indicated, the app reports forest, funnel and cumulative plots, as well as a list of relevant publications. The app is a useful resource, which can be used to *i)* comprehensively retrieve relevant publications, *ii)* explore the literature at an individual researcher’s needs’ level, *iii)* define new hypotheses, *iv)* evaluate publication bias and replicability of findings, and *v)* estimate realistic effect sizes on which to ground future research.

The validity of our conclusions is not limited to the robustness of the models used but grounded on the vast primary evidence included (>200 publications). As a consequence, accidental findings have little weight. Although the methods and approach we adopt are rigorous and reasonably conservative, the quality of the conclusions critically depends on the quality of the studies and data included. From our qualitative bias assessment, the risk for potential bias was lower than previously reported in Neuroscience (30, 54, 55); yet, only ~20% of studies stated being blinded as well as randomized. Furthermore, any meta-analytic dataset is burdened with missing data, due to publication bias or to the preferred investigation of certain factors over others (56). Our models did display evidence of publication bias, yet they were robust to several corrections and sensitivity analyses. Although we cannot fully exclude that the above-mentioned limitations may affect the outcome, it is unlikely that the conclusions drawn would be substantially impacted. Nevertheless, we have attempted to address these methodological issues as comprehensively as possible in our analysis.

### 5.2 Considerations on ELA models

The behavioral changes we report are presumably a convergent phenotype of distinct, model-dependent, underlying biological mechanisms. An organism’s development is not linear nor simultaneous for every component, but it occurs in critical periods (57). For example, postnatal day (P)2-P5 is a sensitive period for the maturation of the adrenal glands (58), P9 for prepulse inhibition (59), and ~P10 for adrenal responsiveness (60). Furthermore, higher cognitive functions develop as multistage processes of sequential nature (57). Accordingly, ELA may particularly disrupt the development of competences whose critical period is active during the time of stress, thereby heightening the variability of the ELA phenotype.

Evidence supporting these notions derives from studies using a single 24h maternal deprivation paradigm, which show a persistent yet paradoxical hypo- and hyper-responsiveness of juvenile ACTH if deprivation occurred at P3 or P11 respectively (61). Thus, while meta-analyses may serve to discern patterns among vast amounts of studies, exploratory studies experimentally dissecting components of ELA in rodents remain indispensable for addressing the underlying mechanisms of action of ELA to the brain (for example: (62, 63)).

#### 5.2.1 Suggestions for future ELA research

Given that the criteria for construct and face validity of ELA models have been met (64), our results provide a practical framework where researchers can anticipate the ELA effect on cognitive outcomes and/or build their own ELA model accordingly. Our exploratory analysis gives insights in the suitability of the models and tests to choose, depending on the question.

Based on this analysis, we tentatively conclude that *i)* rats seem overall more sensitive to ELA-induced changes than mice. Moreover, *ii)* elements such as transporting pregnant dams appear to amplify the effects of ELA. Such stressful life events may have substantial impact on the system, in line with the multiple-hit theory (23). As evident from Figure 3, *iii)* a duration of ~10 days ELA produced the most robust phenotype. Finally, *iv)* the limited bedding and nesting (LBN) model produced the largest effect sizes when compared to separation/deprivation models. Given this reliability, in combination with the feasibility and translational validity, LBN seems an influential paradigm to investigate the mechanisms of chronic stress early in life (39, 65).

To reduce variability and improve comparability across studies, ELA should be preferably applied with consistent protocols (S1.5), unless manipulation of particular aspect(s) of the model is under investigation. Obviously, the importance of individual variation is a factor that should not be overlooked. In our analysis, the paradigm of licking-and-grooming – which is not experimentally induced but based on natural variation in maternal care – consistently evoked the largest effect sizes, although these were based on fewer publications than the other models.

### 5.3 Translational Potential

ELA is one of the most consistent environmental risk factors for the development of psychopathology (2). Although the effects of ELA on the brain can be adaptive, they may evolve into dysfunctional elements in genetically predisposed individuals (2). Behavioral performance in specific cognitive domains seems to be a relevant intermediate phenotype (10), as it may mediate the effects of ELA on psychopathology. For example, in post-traumatic stress disorder, enhanced memory of stressful events becomes pathological after a later-life trauma (8).

In humans, the concept of ELA is extremely varied. Even when considering solely maltreatment, this can be characterized by repeated or sustained episodes of various forms of neglect and abuse (66). Furthermore, the environmental variation is intertwined with socio-economic status, complex relations (e.g. family, neighborhoods, peers, school), and intergenerational transmissions (66). Rodent paradigms do not capture the complexity of human ELA, but they can model specific *aspects* of the human variability in a well-controlled setting. For example, LBN is based on the erratic and unpredictability of maternal care (39, 65, 67) that has been established as a hallmark in childhood abuse situations (68). Similarly, cognitive performance (e.g. memory after stressful learning) can be modelled in rodents, albeit with clear restraints: the tasks are obviously different, should be interpreted in relation to the animal’s normal behavior, and cannot investigate a range of outcomes such as verbal abilities, critical for social interaction and psychopathology (69), also in relation to ELA (70).

Explaining how ELA increases psychopathology risk requires the understanding of its complex interplay with other susceptibility/resilience factors, such as genetic background and later life stressors (71). This mechanistic investigation is difficult to achieve in humans, where limited material, difficulty of prospective and longitudinal designs, complexity and lack of control over the environment and genetic variation hamper causal inferences of ELA to later life cognitive performance. To this end, animal studies can be of considerable added value (39).

An interesting issue in evaluating the translational potential of ELA rodent models is sex differences. In our analysis, males showed larger effect sizes (albeit in the same direction) than females to the effects of ELA on all outcomes, thereby confirming previous preclinical literature (72). Conversely, in clinical populations, females appear more sensitive to childhood trauma as well as to the development of stress-related psychopathologies (39), although sex differences depend on the type of disorder (73). A plausible biological explanation for this discrepancy is the developmental timing during which stress occurs. Although humans and rodents are altricial species, the brain of newborn rats corresponds roughly to 23/24-week old human fetuses (74). Interestingly, the sensitivity to adversities in the last trimester of gestation in humans has been suggested to affect males more than females (75). Experimentally manipulating the timing of ELA exposure may further elucidate ‘female’ stress-sensitive periods. It therefore remains to be established whether the effects of ELA on cognitive domains are truly different between sexes. Our analyses suggest that the effects may not be sexually dysmorphic in nature but may result from the experimental designs used. For example, ELA models and behavioral tests were originally developed for males: maternal care shows clear sex-specific differences (76, 77), and females perform poorly in behavior tests such as object recognition and object-in-location (39, 40). Consequently these paradigms may not be sensitive enough for a female’s phenotype. Indeed, the recorded effects were in the same direction across sexes, and MetaForest attributed to sex a relatively modest variable importance. Our results showcase the necessity to study sex as a biological variable (75, 78), which requires the development of tests and models that are female-specific. This step is required for a more meaningful comparison between rodent and humans, and a delineation of the underlying sex-dependent mechanisms of ELA.

Despite these drawbacks, our meta-analysis confirms and importantly extends standing hypotheses on ELA based on exploratory studies. To aid future investigations in this field, we provide a online tool to evaluate existing literature and direct the experimental design of new studies.

## 6 Acknowledgements

This work was supported by the Consortium on Individual Development (CID), which is funded through the Gravitation program of the Dutch Ministry of Education, Culture, and Science and Netherlands Organization for Scientific Research (NWO grant number 024.001.003). R.A.S. was supported Netherlands Organization for Scientific Research (NWO Veni grant 863.13.02).

We would like to thank Lara Oblak for contributing to the extraction of statistical information and Ruth Damsteegt for discussions on behavioral interpretation. We gratefully acknowledge Prof. Herbert Hoijtink for feedback on the methods, and Prof. Rien van IJzendoorn and Prof. Marianne Bakermans for helpful discussions.

