## Supplemental Material for "The behavioral phenotype of early life adversity: a 3-level meta-analysis of preclinical studies"

### 1 Supplemental methods

#### 1.1 Definitions

The table below reports an explanation/definition of (technical) terms used throughout the manuscript.

| Terms | Definitions and assumptions |
|---|---|
| Behavioral test | Experimental method to measure behavior in a standardized manner |
| Outcome variables | Outcomes reported for each behavioral test.<br>A complete list of included behavioral tests and related variables can be found in S1.3. |
| Individual comparison | Each effect size measured between a control and an experimental group with a history of ELA.<br>Often, multiple outcome variables were measured and reported for each behavioral test performed. In an attempt to limit hierarchy of the data, we rated *a priori* how well a variable described the behavioral domain that the test aimed to operationalize. If multiple variables were reported, we selected only the one with the highest rating. It follows that each behavioral test is represented in the dataset by only one individual comparison. Rating of the variables for each behavior test can be found in S1.3. |
| Experiment | Ensemble of individual comparisons (each representing a different behavioral test) from the same groups of animals.<br>Individual comparisons within the same experiment were considered *dependent* on each other. Any publication can report multiple experiments. Individual comparisons from different experiments within the same publication are considered *independent* of each other as they derive from different animals.<br>If a publication did not mention that different cohorts of animals were |

| | |
|---|---|
| | used, we assumed that all behavioral tests and related outcomes were performed in the same animals and therefore belonged to the same experiment. |
| Nest | Unit of aggregation in the multi-level model. Here, it corresponds to the “experiment” level as individual comparisons within the same experiment derive from the same animals and are therefore *dependent* on each other. |

#### 1.2 Search string

##### 1.2.1 PubMed

(“early life stress”[tiab] OR “ELS”[tiab] OR “early life adversity”[tiab] OR “early life adversities”[tiab] OR “early life adversity*” OR "early stress”[tiab] OR “neonatal stress”[tiab] OR “postnatal stress”[tiab] OR “perinatal stress”[tiab] OR “neonatally stressed”[tiab] OR “early adverse experience”[tiab] OR “perinatally stressed”[tiab] OR “early adverse experiences”[tiab] OR “postnatal manipulation”[tiab] OR “postnatal manipulations”[tiab] OR “perinatal manipulation”[tiab] OR “perinatal manipulations”[tiab] OR “maternal separation”[tiab] OR “maternal deprivation”[tiab] OR “maternal care”[tiab] OR “isolation”[tiab] OR “limited bedding”[tiab] OR “limited nesting”[tiab] OR “limited material”[tiab] OR licking and grooming”[tiab] OR “licking-grooming”[tiab] OR “licking/grooming”[tiab])

AND

(“murine”[tiab] OR “rodentia”[tiab] OR “rodent”[Tiab] OR “rodents”[Tiab] OR “rodentia”[tiab] OR mus[Tiab] OR murinae[Tiab] OR muridae[Tiab] OR “mice”[MeSH Terms] OR “mice”[tiab] OR “mouse”[tiab] OR “rats”[MeSH Terms] OR “rat”[tiab] OR “rats”[tiab])

AND

(“Behavior, Animal”[Mesh] OR “behaviour”[tiab] OR “behavior”[tiab] OR “behaviours”[tiab] OR “behaviors”[tiab] OR “behav*”[tiab] OR “behavioural test”[tiab] OR “behavioural tests”[tiab] OR “behavioral test”[tiab] OR “behavioral tests”[tiab] OR “test, behavioral”[tiab] OR “test, behavioural”[tiab] OR “tests, behavioral”[tiab] OR “tests,behavioural”[tiab] OR “anxiety”[tiab] OR “fear”[tiab] OR “anxiety/fear”[tiab] OR “anxiety-fear”[tiab] OR “emotional learning”[tiab] OR “non-stressful learning”[tiab] OR “non stressful learning”[tiab] OR “social behaviour”[tiab] OR “social behavior”[tiab] OR “sexual behaviour”[tiab] OR “sexual behavior”[tiab] OR “radial arm”[tiab] OR “T maze”[tiab] OR “Ymaze”[tiab] OR “what where which task”[tiab] OR “what-where-which task”[tiab] OR “object in location”[tiab] OR “object in context”[tiab] OR “object recognition”[tiab] OR “object discrimination”[tiab] OR “barnes maze”[tiab] OR “holeboard”[tiab] OR “circular maze”[tiab] OR “Morris water maze”[tiab] OR “spontaneous alteration task”[tiab] OR “maze learning”[tiab] OR “active avoidance”[tiab] OR “spring test”[tiab] OR “inhibitory avoidance”[tiab] OR “passive avoidance”[tiab] OR “defensive withdrawal”[tiab] OR “fear conditioning”[tiab] OR “cat box”[tiab] OR “elevated plus maze”[tiab] OR “EPM”[tiab] OR “cross maze”[tiab] OR “open field”[tiab] OR “concentric square field test”[tiab] OR “place preference”[tiab] OR “place avoidance”[tiab] OR “light/dark test”[tiab] OR “light dark test”[tiab] OR “light-dark test”[tiab] OR “light/dark box”[tiab] OR “light dark box”[tiab] OR “light-dark box”[tiab] OR “object exploration”[tiab] OR “square field test”[tiab] OR “shuttle box”[tiab] OR “social interaction”[tiab] OR “three chambers”[tiab] OR “3 chambers”[tiab] OR “three chamber”[tiab] OR “3 chamber”[tiab] OR “1 chamber”[tiab] OR “one chamber”[tiab] OR “emotional witness stress”[tiab] OR “social play”[tiab] OR “social approach test”[tiab] OR “social encounter test”[tiab] OR “social interaction test”[tiab] OR “social preference test”[tiab] OR “social learning”[tiab] OR “social preference”[tiab] OR “social hierarchy”[tiab] OR “dominance”[tiab] OR “tube test”[tiab] OR “resident test” [tiab] OR “intruder test”[tiab] OR “resident intruder test”[tiab] OR “resident/intruder test”[tiab] OR “resident-intruder test”[tiab] OR “competitive behaviour”[tiab] OR “competitive behaviour”[tiab] OR “play fighting behaviour”[tiab] OR “play fighting behaviour”[tiab] OR “play-fighting behaviour”[tiab] OR “play-fighting behavior”[tiab] OR “play/fighting behaviour”[tiab] OR “play/fighting behavior”[tiab])

##### 1.2.2 WebOfScience

“early life stress” OR “ELS” OR “early life adversity” OR “early life adversities” OR “early life adversity*” OR "early stress” OR “neonatal stress” OR “postnatal stress” OR “perinatal stress” OR “neonatally stressed” OR “early adverse experience” OR “perinatally stressed” OR “early adverse experiences” OR “postnatal manipulation” OR “postnatal manipulations” OR “perinatal manipulation” OR “perinatal manipulations” OR “maternal separation” OR “maternal deprivation” OR “maternal care” OR “isolation” OR “limited bedding” OR “limited nesting” OR “limited material” OR “licking and grooming” OR “licking-grooming” OR “licking/grooming”

AND

“murine” OR “rodentia” OR “rodent” OR “rodents” OR “rodentia” OR mus OR murinae OR muridae OR “mice” OR “mouse” OR “rat” OR “rats”

AND

“behaviour” OR “behavior” OR “behaviours” OR “behaviors” OR “behav*” OR “behavioural test” OR “behavioural tests” OR “behavioral test” OR “behavioral tests” OR “test, behavioral” OR “test, behavioural” OR “tests, behavioral” OR “tests,behavioural” OR “anxiety” OR “fear” OR “anxiety/fear” OR “anxiety-fear” OR “emotional learning” OR “non-stressful learning” OR “non stressful learning” OR “social behaviour” OR “social behavior” OR “sexual behaviour” OR “sexual behavior” OR “radial arm” OR “T maze” OR “Ymaze” OR “what where which task” OR “what-where-which task” OR “object in location” OR “object in context” OR “object recognition” OR “object discrimination” OR “barnes maze” OR “holeboard” OR “circular maze” OR “Morris water maze” OR “spontaneous alteration task” OR “maze learning” OR “active avoidance” OR “spring test” OR “inhibitory avoidance” OR “passive avoidance” OR “defensive withdrawal” OR “fear conditioning” OR “elevated plus maze” OR “EPM” OR “cross maze” OR “open field” OR “concentric square field test” OR “place preference” OR “place avoidance” OR “light/dark test” OR “light dark test” OR “light-dark test” OR “light/dark box” OR “light dark box” OR “light-dark box” OR “object exploration” OR “square field test” OR “shuttle box” OR “social interaction” OR “three chambers” OR “3 chambers” OR “three chamber” OR “3 chamber” OR “1 chamber” OR “one chamber” OR “emotional witness stress” OR “social play” OR “social approach test” OR “social encounter test” OR “social interaction test” OR “social preference test” OR “social learning” OR “social preference” OR “social hierarchy” OR “dominance” OR “tube test” OR “resident test” OR “intruder test” OR “resident intruder test” OR “resident/intruder test” OR “resident-intruder test” OR “competitive behaviour” OR “competitive behaviour” OR “play fighting behaviour” OR “play fighting behaviour” OR “play-fighting behaviour” OR “play-fighting behavior” OR “play/fighting behaviour” OR “play/fighting behavior”

#### 1.3 CLASSIFICATION OF BEHAVIORAL TESTS IN BEHAVIORAL DOMAINS

Prior the beginning of the study, four experts (JK, RvdV, MJ & Ruth Damsteegt) were consulted for the selection of tests and related outcomes, as well as for their classification in behavioral domains. Overall, we aimed to extract the variable for each test that best represented the described domain. However, often the most reported outcome for a certain test is not necessarily the best at representing the categorized domain. For example, in the anxiety test "elevated plus maze", the most reported outcome is "time spent in open arms". Arguably "time spent in the closed arms" is a more direct measure of anxiety: the more anxious the animal, the more the time spent in closed arms. In such circumstances, if a paper reported both outcomes, we extracted the most common (in this case, time spent in the open arms), in the intent to avoid unnecessary heterogeneity. The experts agreed on which variables for each test best expressed the categorized behavioral domain (e.g. anxiety, memory), and ranked them based on their importance.

Table legend: **Importance** = ranking for variable selection (for details see above); **multiplication effect size for model** = effect sizes were multiplied whenever necessary by -1 so that an increase in Hedge's G would indicate an increase in anxiety, improved memory after stressful learning, impaired memory after neutral learning and decreased social behavior (Section **Error! Reference source not found.**). † = Tests included in the systematic review, but not in the confirmatory analysis. Specific inclusion/exclusion criteria for these tests are specified in the footnote below.

| Behavioral test | Outcomes reported | Importance | Multiplication effect size for model | Comments |
|---|---|---|---|---|
| **Anxiety** | | | | |
| Defensive withdrawal | Time spent in the center | 1 | -1 | |
| | Time spent in the tube | 2 | 1 | |
| | Latency to exit the tube | 3 | 1 | |
| Elevated zero maze | Time spent in open arms | 1 | -1 | |
| | Time spent in closed arms | 2 | 1 | |
| | entriesOpen | 3 | -1 | |
| Elevated plus maze | Time spent in open arms | 1 | -1 | |

| | | | | |
|---|---|---|---|---|
| EPM | Amount entries in open arms | 2 | -1 | |
| Fear conditioning | Amount of time spent freezing | 1 | 1 | After first footshock (no retention) |
| Forced swim test | Time spent immobile | 1 | 1 | during first exposure |
| | Time spent struggling | 2 | -1 | during first exposure |
| Light/Dark Box | Time spent in the dark compartment | 1 | 1 | |
| | Time spent in the light compartment | 2 | -1 | |
| | Latency to enter the dark compartment | 3 | -1 | animal starts test in light compartment |
| Novelty-induced reduction of feeding/drinking | Latency to feed | 1 | 1 | |
| | Time spent in the center | 2 | -1 | |
| | Time spent feeding | 3 | -1 | |
| Open field * | Time spent in the center | 1 | -1 | |
| | Time spent in the periphery | 2 | 1 | |
| | Distance moved in the center | 3 | -1 | |
| | Latency to enter the center | 4 | 1 | animal starts test in the periphery |
| | Amount entries in the center | 5 | -1 | |
| Tail suspension test | Amount of time spent immobile | 1 | 1 | |
| **Memory after stressful learning** | | | | |
| Fear Conditioning | Amount of time spent freezing | 1 | 1 | At re-exposure (retention time scored in separate variable). Fear can also be "social" |
| Forced swim test | Time spent immobile | 1 | 1 | At re-exposure (retention time scored in separate variable) |
| | Latency to immobility | 2 | -1 | At re-exposure (retention time scored in separate variable) |
| | Distance moved | 3 | -1 | At re-exposure (retention time scored in separate variable) |
| | Frequency immobility scored | 4 | 1 | At re-exposure (retention time scored in separate variable) |
| Morris water Maze (water temperature | Time spent in target quadrant | 1 | 1 | |
| | Distance swum in target quadrant | 2 | 1 | |

| | | | | |
|---|---|---|---|---|
| <24°C)* | Latency to find platform | 3 | -1 | If probe trial not present* |
| Shuttle box | Amount of avoidance responses | 1 | 1 | |
| | Latency to avoid | 2 | -1 | |
| | Latency to enter avoidance compartment | 2 | -1 | Same as latency to avoid |
| **Memory after neutral learning** | | | | |
| Morris water Maze (water temperature >26°C)* | Time spent in target quadrant | 1 | 1 | |
| | Distance swum in target quadrant | 2 | 1 | |
| | Latency to find platform | 3 | -1 | If probe trial not present* |
| Object in context | Discrimination index | 1 | -1 | |
| Object in location | Discrimination index | 1 | -1 | |
| | Time spent with novel object | 2 | -1 | |
| Object recognition | Discrimination intex | 1 | -1 | |
| | Time spent with novel object | 2 | -1 | |
| | Ratio time spent novel / familiar object | 3 | -1 | |
| Social recognition | Discrimination index | 1 | -1 | novel vs familiar animal |
| | Time spent with novel animal | 2 | -1 | novel vs familiar animal |
| Temporal order task | Discrimination index | 1 | -1 | |
| T maze | Time spent in novel arm | 1 | -1 | |
| Y maze | Time spent in novel arm | 1 | -1 | |
| **Social behavior** | | | | |
| Resident intruder test | Time spent in aggressive behavior | 1 | 1 | |
| | Tiime spent attacking | 2 | 1 | |
| | Latency to first aggression | 3 | -1 | |
| | Latency to first attack | 4 | -1 | |
| | Total amount of aggressive behavior | 5 | 1 | |

| | | | | |
|---|---|---|---|---|
| | Time spent in offensive posture | 6 | 1 | |
| Social interaction | Time spent in social interaction | 1 | -1 | |
| | Amount of social interaction | 2 | -1 | |
| Social open field | Time spent in social proximity | 1 | -1 | |
| Social play | Time spent in social interaction | 1 | -1 | |
| Social preference | Preference index | 1 | -1 | animal vs inanimate object |
| | Time spent in social interaction | 2 | -1 | |
| | Time spent in social tube | 3 | -1 | |
| **Other †** | | | | |
| 8-arms radial maze | Errors in working memory | | | |
| T maze | Alternation | | | |
| Y maze | Alternation | | | |
| Step down inhibitory avoidance | Latency to step down | | | at retest |
| Morris Water Maze * | | | | If water temperature between 24°C and 26°C |

* = Inclusion/exclusion criteria for specific tests:

- Open field is included only if test length <15 min. If test length >15min, the test is considered a measure of locomotor activity and not anxiety.
- In the open field, "amount of crossings" is considered a measure of locomotor activity and not anxiety. Tests reporting this as the only measure are not included.
- The Morris Water Maze test is considered stressful if water temperature <24°C, non-stressful if water temperature >26°C. Water "at room temperature" was considered as 24°C. If water temperature was between 24°C and 26°C, it was considered not classifiable in either the non-stressful or stressful domain. Nonetheless, these comparisons were included in the exploratory part.

- Working memory is excluded from meta-analysis due to controversial domain categorization in memory after stressful / non-stressful learning. Nonetheless, it is included in the systematic review as it provides information about memory retention and repetition.
- Step down-inhibitory avoidance is excluded from the meta-analysis because it is questionable whether for an animal remaining on a platform for a long time would be better or worse than an inescapable footshock. To the experts, it was therefore controversial to define the directionality of the effect. Nonetheless, we include this test in the systematic review. Furthermore, we ran a sensitivity analysis by including step down-inhibitory avoidance as part of the memory after stressful learning domain and verified that the interpretation did not change.

#### 1.4 DEFINITION OF MULTIPLE HITS

Prior to the beginning of the study, we defined elements that would constitute “multiple hits”(1). Although this would ideally be a continuous variable (e.g. severity), we solely categorize its presence/absence due to the complexity and subjectivity of the classification. Animals were considered in the “multiple hits” group if they had one of the following (in addition to ELA):

| Considered multiple hits | Not considered multiple hits |
|---|---|
| Stressful behavioral test performed previously (e.g. FST, fear conditioning) | Intragastric saline |
| Footshocks | Saline injections |
| Chronic (mild) unpredictable stress | Vaginal smears |
| Chronic constant light | Daily handling by experimenter |
| Chronic restraint | |
| Chronic individual housing | |
| Vaginal balloon distention | |
| Cannula implantation, mock surgeries, blood sampling, isofluorane anaesthesia | |
| Dams transported pregnant | |
| Stress prone strain (BALB/C, wistar Kyoto, DBA) | |

**Note: manipulated genetic background were excluded from the meta-analysis (S1.5) and therefore could not be included in the definition of vulnerability, despite it being an important factor

#### 1.5 INCLUSION/EXCLUSION CRITERIA

Study selection was performed independently by two researchers (VB and JK), who were blinded to the studies' results. The inclusion and exclusion criteria were specified prior to the beginning of the study.

| Criteria | Comments |
|---|---|
| Inclusion | |
| Peer reviewed original publications in English | |
| Mice and rats | |
| ELA starts before P14 | ELA model can extend after P14 |
| ELA as alteration of maternal care(2)<br>• separation of the pup from the mother (maternal separation(3) / deprivation(2))<br>• separation of the pup from mother and siblings (isolation)<br>• limited bedding and nesting(4)<br>• licking and grooming(5) | We define as '*separation'* those models in which the mother was repeatedly separated from the pups (e.g. 3 hours a day for 2 weeks). We define as *deprivation* those models in which the mother was separated once from the pups for a prolonged time (e.g. 1 time 24 hours, or 2 times 12 hours). In other words, the categorization in maternal separation/deprivation depends on the model used and not the naming used in the papers.<br>The separation/deprivation/isolation models are adaptations of Levine's original model. These adaptations are based on the observation that dams often leave the nest to forage for 15-30 min periods(6). For this reason, we consider "adverse" and therefore include only those studies in which the duration of separation/deprivation/isolation time was >1h. |
| Behavioral testing during adult age | Older than 8 weeks but younger than 1 year |
| Behavioral tests among the listed *a priori* | See 1.3 |

| Exclusion | |
|---|---|
| Specific pathogen free animals | |
| Ovariectomized females | |
| Sex not specified | Publication is included if sex is retrieved after contacting the authors |
| Males and females pooled | Publication is included if summary statistics of males and females separately are received after contacting the authors |
| Fasting before behavioral test (unless part of the test itself) | |
| Handling, gentling and communal nesting as ELA models | |
| Maternal separation with early weaning(7) | Early weaning is defined as separation of the pups from the mother at P17.<br>If early weaning is only in the experimental group, the experiment is excluded. If early weaning occurred in both control and experimental group, the study is included and early weaning is considered a factor that could increase vulnerability |
| Handling as control group | |
| Genetic manipulations | |
| Animals bred for high/low anxiety or novelty response or sensitivity/resilience to depression | |
| Animals separated in high/low performance | |
| Administration of any drug or alcohol via any route | e.g. Drug injections before testing, methamphetamine conditioned place preference tests |
| Any manipulation to previous generations | |
| Other * | Inclusion/exclusion criteria specific to certain tests. See footnote of S1.3. |

Choosing the outcomes to include in the meta-analysis is often a non-straightforward task. Despite our intent to be as comprehensive as possible in the definition of inclusion/exclusion criteria *prior* to the beginning of the study, the list was not exhaustive. We therefore added inclusion/exclusion criteria *during* data collection:

- Unless the test required training, the test is included only during the first exposure
- If the data is presented in multiple time bins, we extracted the mean value reported most similar to other papers of the same category. In particular, we selected:
  - The first time bin for anxiety behaviors
  - The last time bin for learning during the Morris Water Maze (intent to be as similar as possible to the probe test)
- Memory extinction and reversal learning were excluded

#### 1.6 Details extraction of statistical information

Effect size was preferably calculated from mean, standard deviation (SD) and amount of animals ($n$) for each group (control and experimental). If only the standard error of the mean (SEM) was reported, SD was calculated as $SEM * \sqrt{n}$. If the number of animals was reported as a range (e.g. 6-8 animals per group), we used the mean of this number (e.g. 7 animals per group). If median and interquartile range (IQR) were reported instead of mean and SD, we assumed normality and considered median = mean and the IQR as SEM/6(8). If total $n$ was provided, $n$ was equally split across groups. If $n$ was not mentioned, could not be calculated from the degrees of freedom nor could be retrieved from the authors, we considered $n$ to be equal to the $n$ average of all other comparisons combined.

If a single control group was used to compare experimental groups in which ELA was induced with different models (e.g. maternal deprivation at P4 vs maternal deprivation and P9(9)), the sample size of the control group was equally divided as control for each experimental group (e.g. n=10 overall in control group becomes n=5 for control of maternal deprivation at P4 and n=5 for control of maternal deprivation at P9)(10).

In our intent to classify as extensively as possible the methodological heterogeneity between different studies, we categorize >40 variables. However, not every publication reported on each of these, giving rise to missing information. Due to model estimation requirements with MetaForest(11), it is necessary to estimate missing values. Following standard practice, we imputed the median of the variable of interest in case of continuous variables, and the most common category in case of categorical variables.

#### 1.7 Sensitivity analysis & Analysis of influential cases

We conducted the following sensitivity analysis:

- Specified *prior* to the analysis:
  - Outlying and influential cases
    - We identified outlying and influential cases according to Viechtbauer & Cheung's definition (12). We qualitatively investigated the identified outlying and influential cases, but we could not identify any specific pattern of characteristics. The identified comparisons were removed and we evaluated the consistency of the results as sensitivity analysis
    - We identified potentially outlying cases also according to Tabachnick and Fidell's definition (13). We conducted a sensitivity analysis by removing them from the analysis and verifying results' consistency.
  - Blinded and randomized studies
    - According to the standards of meta-analysis, we should include only studies which were blinded as well as randomized. However, only a few comparisons had these characteristics. For this reason, we chose to perform the main analysis on the full dataset.
    - To check for the influence of blinding and randomization on the effects sizes estimated in the main analysis, we performed a sensitivity analysis by including a "blinded and randomized" variable as a moderator in our model. We confirmed that this moderator was not significant (**males:** $Q(1) = .316$, $p = .574$; **females:** $Q(1) = 3.263$, $p = .07$).
  - Tests which were only reported by at least 4 publications (including second hit)
  - Risk of potential bias
    - We evaluated whether increase in potential bias corresponded to an increase in effect sizes.

#### 1.8 PUBLICATION BIAS ASSESSMENT DETAILS

Publication bias was assessed with several methods. Although this may seem redundant, this approach was selected to balance out the pros and cons of each method

| Test | Pros | Cons |
|---|---|---|
| Qualitative investigation of funnel plot | Estimated values derive from the built 3-level mixed effect model | Qualitative and not quantitative |
| Egger's regression followed by test for funnel plot asymmetry | Frequently used, quantitative | Does not consider the 3-level design |
| Begg's test | Frequently used, quantitative | Does not consider the 3-level design |
| File drawer analysis with fail and safe test | Addresses file drawer problem (only significant results are published). It provides an estimate of how many studies are necessary to nullify the effect found | Does not consider the 3-level design nor the moderators of the effect |
| Trim and fill | Aims to both identify and correct funnel plot asymmetry. It provides an estimate of the number of missing studies | Does not consider the 3-level design nor the moderators of the effect. Furthermore, it is known to perform poorly when substantial heterogeneity is present. |

#### 1.9 Tuning parameters MetaForest

MetaForest's tuning parameters were selected from a 10-fold cross-validation according to the author's instructions(14). We tested which type of weights (random-effects, fixed-effects or unweighted) provided the best model fit, with how many moderators available at each split (2, 4 or 6), and what was the most appropriate minimum size of the node (2, 4, or 6) to allow for splitting. Root-mean-square error (RMSE) was used to select the optimal model using the smallest value. This led to the selection of the following parameters: uniform weighting metaForest, 4 candidate moderators available at each split, 2 as minimum node size. The estimated residual heterogeneity of the model was tau2 = .46. We investigated the marginal bivariate relationship of each moderator by averaging its effect size over the values of all other moderators. The resulting partial dependency graphs can be obtained with the R code accompanying the text.

### 2 SUPPLEMENTARY RESULTS

#### 2.1 MISSING VALUE DETAILS

We were not able to retrieve information from the following publications:

- 14 manuscripts published before 2008:
    - (15–24)
- Full text of 8 publications was not found (authors contacted):
    - (25–32)
    - It cannot be evaluated whether they were suitable for inclusion
- Authors from 9 manuscripts were contacted but no answer was received:
    - (22, 33–39)

#### 2.2 COMPARISONS EXCLUDED FROM META-ANALYSIS

Below we provide details on comparisons excluded from the meta-analysis due to controversial domain categorization (S1.3). These comparisons were nonetheless analyzed at a systematic review level and are present in the published dataset (https://osf.io/ra947/).

Table legend: $n_{comp}$= amount of comparisons, $n_{exp}$ = amount of experiments from which the comparisons were retrieved, $n_{stud}$ = amount of studies from which the comparisons were retrieved.

| Test | $n_{comp}$ | $n_{exp}$ | $n_{stud}$ | Comments |
|---|---|---|---|---|
| Morris water maze | 4 | 4 | 4 | Water temperature between 24 and 26°C |
| 8 arm radial maze | 2 | 2 | 1 | Working memory |
| T maze | 1 | 1 | 1 | Working memory |
| Y maze | 9 | 7 | 3 | Working memory |
| Step down avoidance | 6 | 6 | 6 | |

#### 2.3 DESCRIPTIVE INFORMATION ON STUDY CHARACTERISTICS

##### 2.3.1 ELA models

Several ELA models are used in the literature to disrupt maternal care. Primarily, these can be distinguished according to the **type** (=which paradigm) and the **timing** (=which postnatal day) of the model. Furthermore, there are specific characteristics within each model that can be altered. These are:

- For separation/deprivation/isolation: the animals are placed in a new cage or remain in the homecage
- For separation/deprivation/isolation: the duration of separation/deprivation/isolation can differ in length
- For separation/deprivation/isolation: the control group can be untouched, animal facility reared, handled <5min, or can derive from a split-litter design
- For separation/isolation: the protocol can be either predictable (every day at roughly the same time) or unpredictable
- For separation/isolation: the protocol occurs during the light/dark phase of the cycle

When considering **timing** as critical periods (model in first-and-second or first-to-third postnatal week): we identified 41 different protocols (322 theoretically possible). In particular, 14 variations of ELA models made up 85.8% of comparisons. This suggests that although there are variations in the protocols, the models are fairly consistent across the literature.

The table below provides descriptive information on the quantity of comparisons ($n_{comp}$) and of relative experiments ($n_{exp}$) across ELA models.

| ELS Model | $n_{comp}$ | $n_{exp}$ |
|---|---|---|
| Maternal separation | 347 | 203 |
| Maternal deprivation | 82 | 49 |
| Isolation | 186 | 104 |
| Limited nesting and bedding | 76 | 34 |
| Licking and grooming | 20 | 12 |

##### 2.3.2 Species and strains

The table below describes the amount of experiments ($n_{exp}$) for each strain used.

| Species | Strain | $n_{exp}$ |
|---|---|---|
| **Mice** | BalbC | 11 |
| | C57Bl/6 | 65 |
| | CD1 | 7 |
| | DBA | 4 |
| | NMRI | 3 |
| | Other | 6 |
| | swissWebster | 1 |
| **Rats** | Lister Hooded | 3 |
| | Long Evans | 19 |
| | Long Evans Hooded | 9 |
| | SpragueDawley | 78 |
| | Wistar | 181 |
| | Wistar Kyoto | 2 |
| | Other | 3 |
| | Not specified | 10 |

##### 2.3.3 Age

The histogram below displays the distribution of age of the animals at the time of testing expressed as postnatal week. We included animals tested for behavior older than 8 weeks of age, but younger than 1 year (S1.5). Although it has been reported that the effects of ELA on behavior (memory in particular) may become more evident in older animals, the amount of comparisons of this age group in our study was not sufficient to further explore this hypothesis.

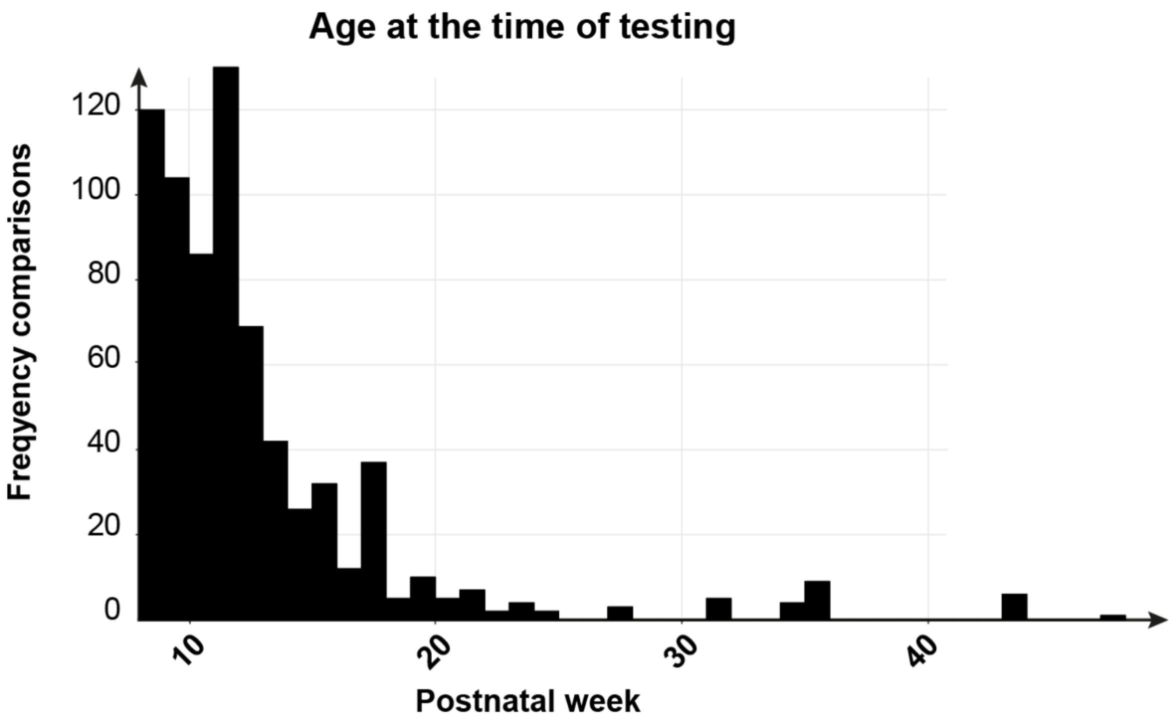

##### 2.3.4 Domains and tests

The tables below display the distribution of comparisons ($n_{comp}$), experiments ($n_{exp}$) and studies ($n_{stud}$) across **A)** sex and domains, **B)** behavioral tests.

| A) | Males | | | Females | | |
|---|---|---|---|---|---|---|
| Domain | $n_{comp}$ | $n_{exp}$ | $n_{stud}$ | $n_{comp}$ | $n_{exp}$ | $n_{stud}$ |
| Anxiety | 262 | 198 | 135 | 95 | 69 | 50 |
| memory after stressful learning | 151 | 136 | 88 | 52 | 45 | 27 |
| Memory after non-stressful learning | 79 | 56 | 45 | 26 | 19 | 17 |
| Social behavior | 40 | 36 | 29 | 6 | 6 | 5 |

| **B)** | **males** | | | **females** | | |
|---|---|---|---|---|---|---|
| **Test** | $n_{comp}$ | $n_{exp}$ | $n_{stud}$ | $n_{comp}$ | $n_{exp}$ | $n_{stud}$ |
| Defensive Withdrawal | 7 | 7 | 4 | 1 | 1 | 1 |
| Elevated Zero Maze | 12 | 11 | 7 | 1 | 1 | 1 |
| Elevated Plus Maze | 105 | 105 | 77 | 38 | 38 | 29 |
| Fear conditioning (anxiety) | 17 | 17 | 12 | 7 | 7 | 7 |

| | | | | | | |
|---|---|---|---|---|---|---|
| Forced Swim Test (anxiety) | 29 | 29 | 19 | 5 | 5 | 4 |
| Light/Dark box | 19 | 19 | 11 | 9 | 9 | 5 |
| Novelty induced-suppression of feeding and drinking | 6 | 6 | 5 | 1 | 1 | 1 |
| Open field | 64 | 64 | 46 | 32 | 32 | 19 |
| Tail suspension test | 3 | 3 | 3 | 1 | 1 | 1 |
| Fear conditioning (stressful learning) | 45 | 37 | 28 | 18 | 13 | 12 |
| Forced swim test (stressful learning) | 52 | 52 | 37 | 17 | 17 | 10 |
| Morris water maze (stressful learning) | 36 | 31 | 25 | 6 | 6 | 5 |
| Shuttle box | 17 | 17 | 6 | 11 | 11 | 5 |
| Social Fear Conditioning | 1 | 1 | 1 | | | |
| Morris water maze (neutral learning) | 2 | 2 | 2 | | | |
| Object in Context | 2 | 2 | 2 | 1 | 1 | 1 |
| Object in Location | 12 | 10 | 9 | 5 | 5 | 5 |
| Object Recognition | 39 | 35 | 31 | 15 | 15 | 14 |
| Social Recognition | 15 | 13 | 9 | 3 | 3 | 2 |
| Temporal Order Task | 4 | 4 | 4 | | | |
| Y Maze (neutral memory) | 6 | 6 | 5 | 2 | 2 | 2 |
| Resident intruder test | 10 | 8 | 7 | | | |
| Social Interaction | 23 | 23 | 18 | 6 | 6 | 5 |
| Social Open Field | 2 | 2 | 1 | | | |
| Social Play | 1 | 1 | 1 | | | |
| Social Preference | 4 | 4 | 3 | | | |
| Morris Water Maze (excl) | 4 | 4 | 4 | | | |
| Radial Maze 8 Arm | 1 | 1 | 1 | 1 | 1 | 1 |
| Step Down Avoidance | 6 | 6 | 6 | | | |
| T Maze | 1 | 1 | 1 | | | |
| Y Maze (excl) | 7 | 7 | 3 | 2 | 2 | 1 |

#### 2.4 Risk of Bias Assessment

Risk of bias assessment was performed according to SYRCLE guidelines, and by distinguishing between bias at an experiment- or study- level. No publication reported information on all SYRCLE potential bias items. Overall, "not specified" was the most common score (54.8%). In 44.8% of the cases, measures to prevent bias were reported. This includes computerized approaches. 41 studies yielding a total of 145 comparisons reported being blinded as well as randomized. For sensitivity analysis, amount of potential bias was operationalized by summing the risk of bias of each item according to the definition: "yes" = 0, "unclear" = 0.5, "no" = 1. Computerized approaches were considered as "0" bias. This produced a continuous variable between 0 (no risk bias) and 10 (maximum risk of bias).

Figure legend: N = Bias was not prevented; NS = it was not specified whether measure to prevent bias were applied; C = computerized approach; Y = measures to prevent bias were used. Percentages on the x axis are in terms of either experiments or publications (right y axis).

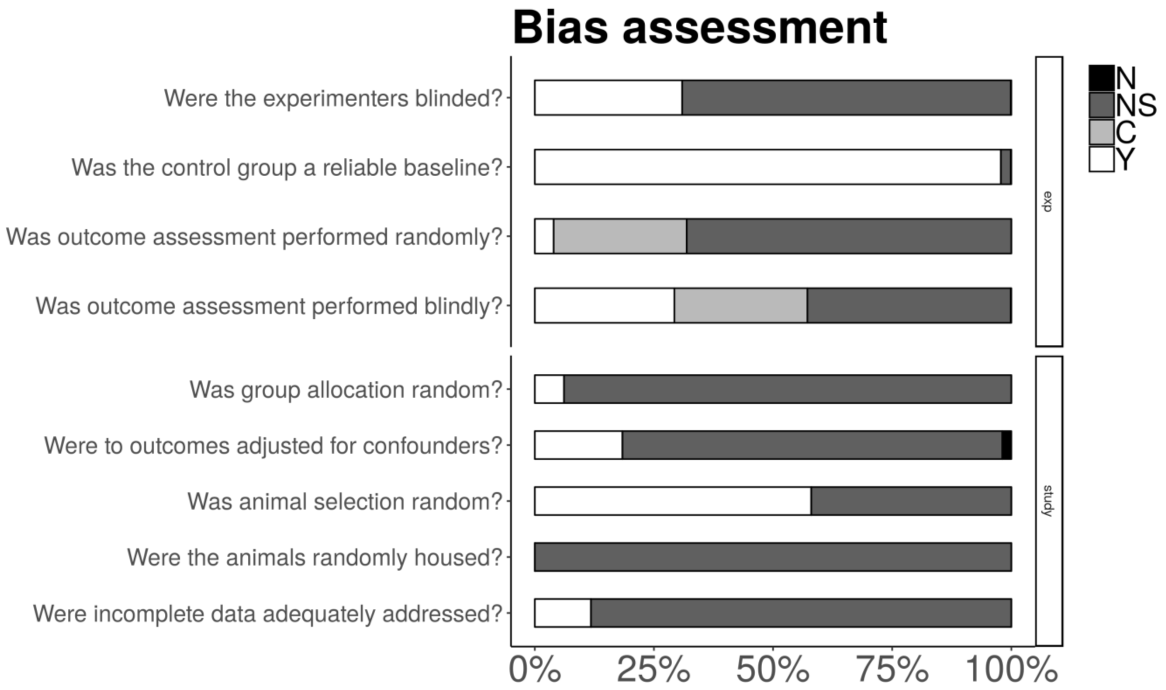

#### 2.5 RESULTS AT A SYSTEMATIC REVIEW LEVEL

On a systematic review level, we evaluated the directionality of the effects of ELS on each behavioral test used. These are expressed as decrease, ns = not significant, increase, and notApplicable = it could not be deduced directly from the data reported. “Increase” should be interpreted as an enhancement of the behavior reported (more anxious, more memory, more social behavior). For example, “increase” in the elevated plus maze signifies that the animals were more anxious. This could mean that they spent less time in or entered fewer times into the open arms. The figures below represent data for each behavior test at a systematic review level in **A)** anxiety, **B)** memory after stressful learning, **C)** memory after non-stressful learning, **D)** social behavior, and **E)** tests not included in the meta-analysis.

##### A) Anxiety

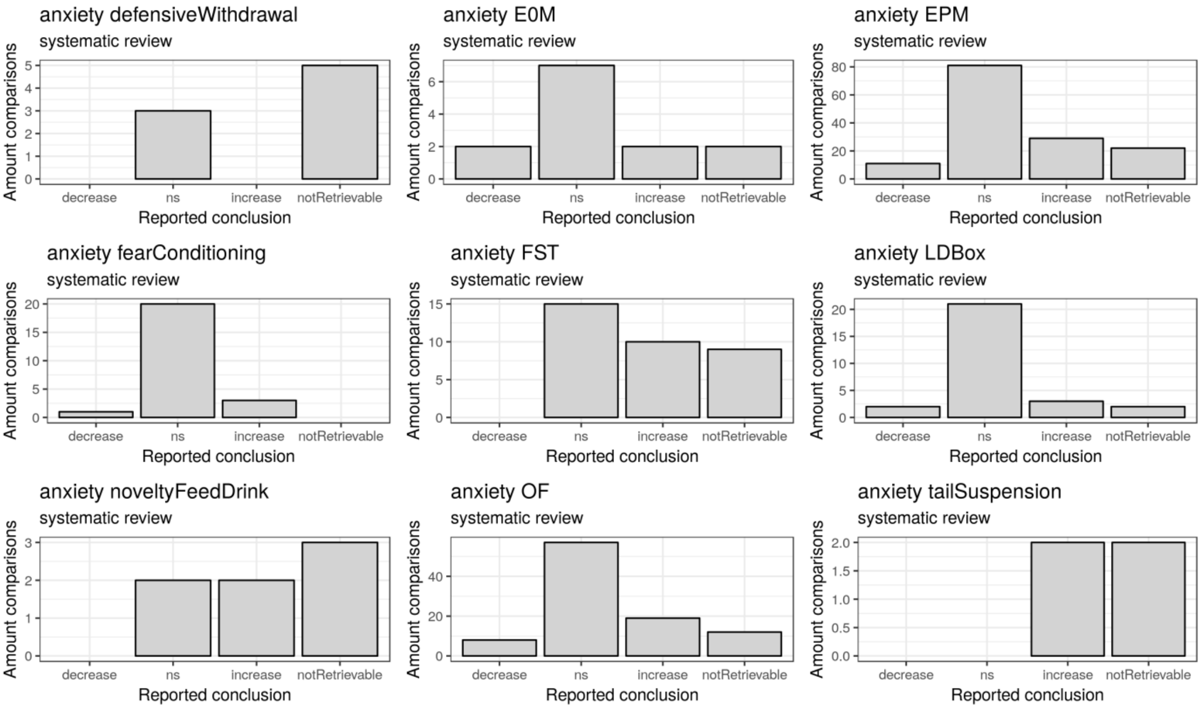

#### B) Memory after stressful learning

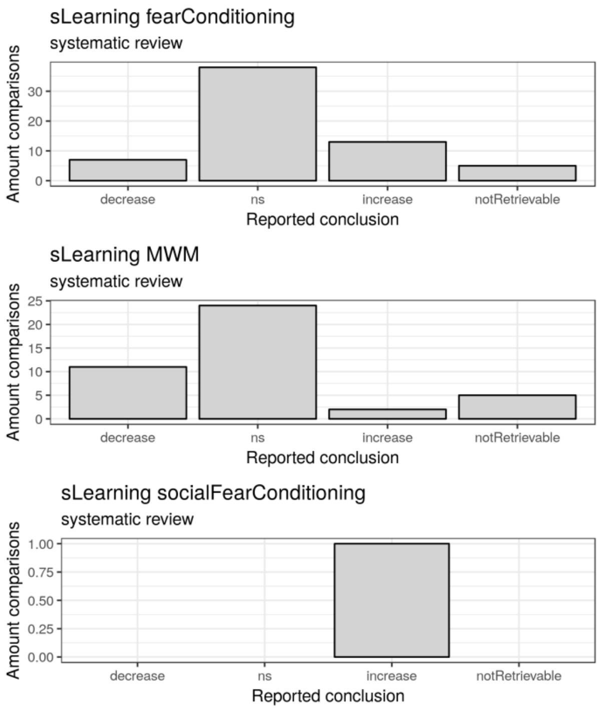

#### C) Memory after non-stressful learning

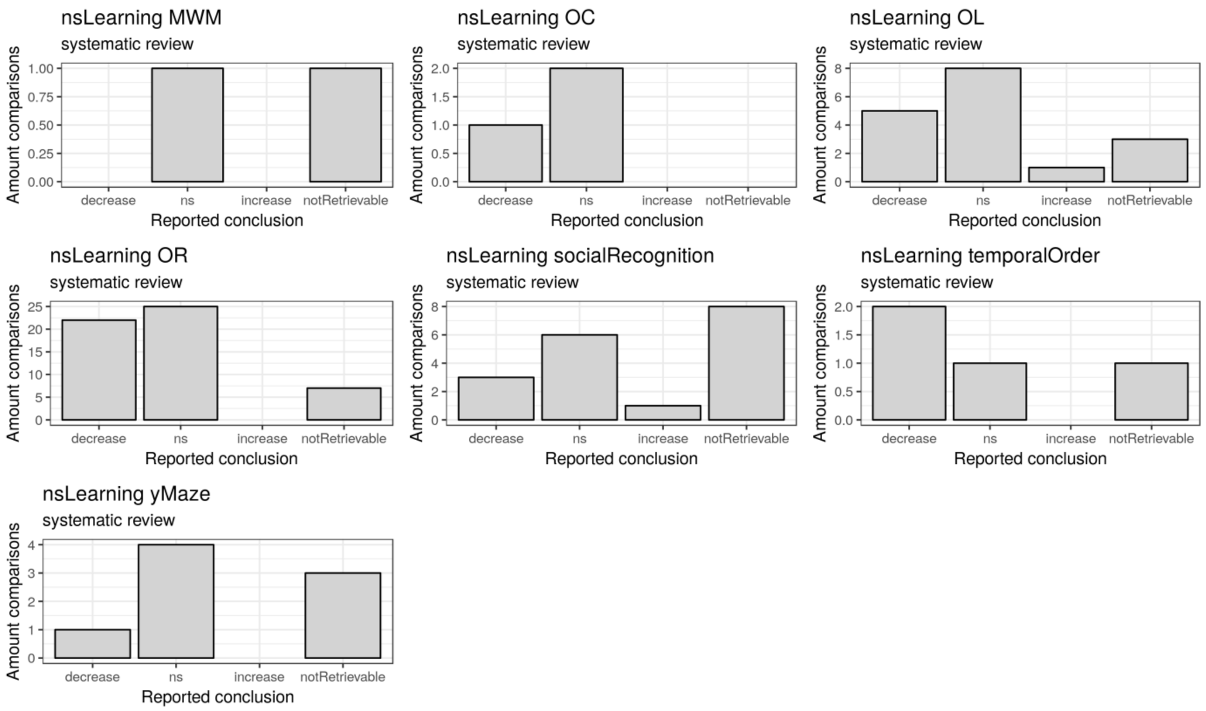

#### D) Social behavior

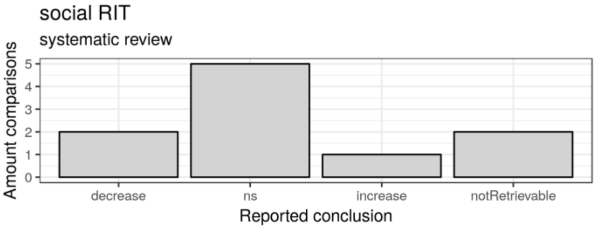

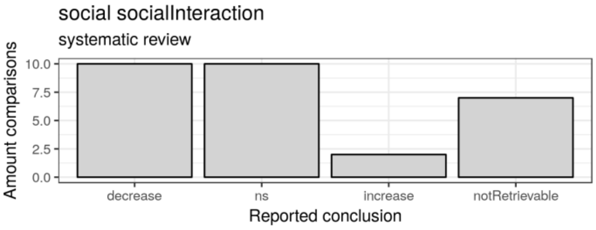

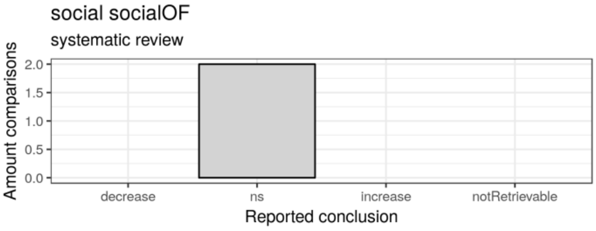

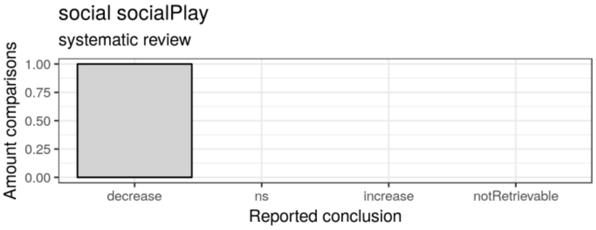

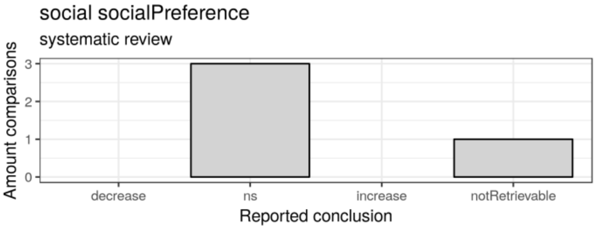

#### E) Not included in the meta-analysis

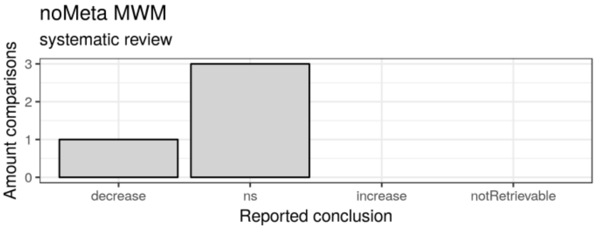

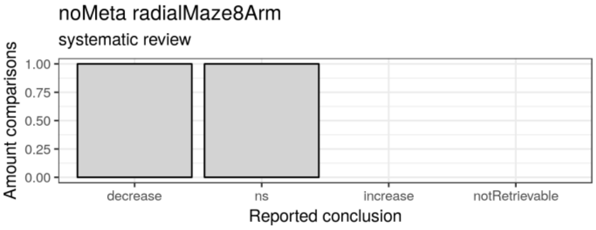

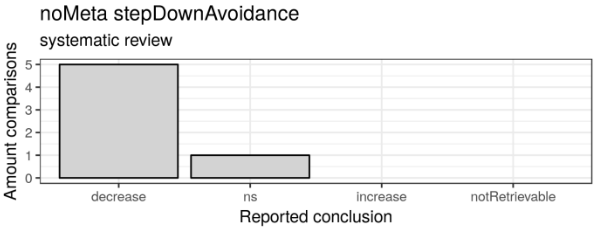

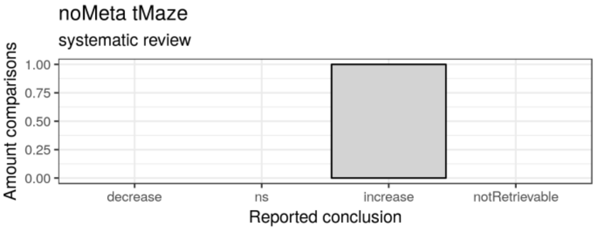

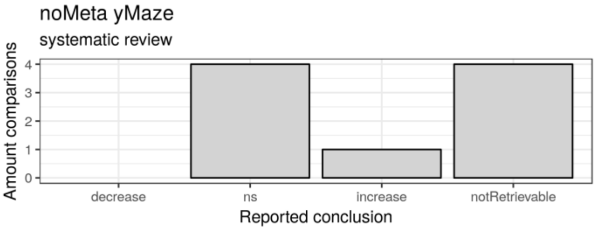

#### 2.6 STATISTICS MAIN RESULTS: MALES

The table below summarizes the results of the hypotheses-testing analysis in males.

**Table legend**: ci.lb = lower boundary confidence interval; ci.ub = upper boundary confidence interval; effectsize = estimated Hedge's G; se = standard error of the estimated Hedge's G; z-value = z-value of the test; p-value = uncorrected p-value of the test; p-value_bonf = corrected p-value for family-wise comparison with Bonferroni; _Corr = refers to effectsize, ci.lb and ci.ub which have been flipped to ease interpretation (Method Section); sLearning = memory after stressful learning; nsLearning = memory after non-stessful learning; domainHit = statistics of experiments with multiple hits vs experiments without; domainNo = only experiments without multiple hits; domainYes = only experiments with multiple hits.

| | test | ci.lb | ci.ub | effectsize | se | Zvalue | Pvalue | Pvalue_bonf | effectsize_Corr | ci.lb_Corr | ci.ub_Corr |
|---|---|---|---|---|---|---|---|---|---|---|---|
| **Domains: main effects** | | | | | | | | | | | |
| | anxiety | 0.1649 | 0.3911 | 0.278 | 0.0577 | 4.8194 | 0 | 0 | 0.278 | 0.1649 | 0.3911 |
| | sLearning | 0.1409 | 0.4255 | 0.2832 | 0.0726 | 3.8997 | 0.0001 | 0.0004 | 0.2832 | 0.1409 | 0.4255 |
| | nsLearning | 0.3953 | 0.7923 | 0.5938 | 0.1013 | 5.8604 | 0 | 0 | -0.5938 | -0.7923 | -0.3953 |
| | social | 0.3477 | 0.8801 | 0.6139 | 0.1358 | 4.5211 | 0 | 0 | -0.6139 | -0.8801 | -0.3477 |
| **Increased vulnerability: main effect** | | | | | | | | | | | |
| | hit | 0.018 | 0.4264 | 0.2222 | 0.1042 | 2.1315 | 0.0331 | 0.0331 | 0.2222 | 0.018 | 0.4264 |
| **Effect increased vulnerability for each domain** | | | | | | | | | | | |
| | anxietyHit | -0.0619 | 0.3811 | 0.1596 | 0.113 | 1.4123 | 0.1579 | 0.6316 | 0.1596 | -0.0619 | 0.3811 |
| | sLearningHit | -0.095 | 0.4676 | 0.1863 | 0.1435 | 1.2989 | 0.194 | 0.776 | 0.1863 | -0.095 | 0.4676 |
| | nsLearningHit | 0.0395 | 0.8309 | 0.4352 | 0.2019 | 2.1556 | 0.0311 | 0.1244 | -0.4352 | -0.8309 | -0.0395 |
| | socialHit | -0.4228 | 0.638 | 0.1076 | 0.2706 | 0.3975 | 0.691 | 1 | -0.1076 | -0.638 | 0.4228 |
| **Posthocs** | | | | | | | | | | | |
| | anxietyNo | 0.0477 | 0.3487 | 0.1982 | 0.0768 | 2.5808 | 0.0099 | 0.0792 | 0.1982 | 0.0477 | 0.3487 |
| | sLearningNo | 0.0031 | 0.3771 | 0.1901 | 0.0954 | 1.9921 | 0.0464 | 0.3712 | 0.1901 | 0.0031 | 0.3771 |
| | nsLearningNo | 0.1 | 0.6524 | 0.3762 | 0.1409 | 2.6699 | 0.0076 | 0.0608 | -0.3762 | -0.6524 | -0.1 |
| | socialNo | 0.2084 | 0.912 | 0.5602 | 0.1795 | 3.1208 | 0.0018 | 0.0144 | -0.5602 | -0.912 | -0.2084 |
| | anxietyYes | 0.1922 | 0.5234 | 0.3578 | 0.0845 | 4.2349 | 0 | 0 | 0.3578 | 0.1922 | 0.5234 |
| | sLearningYes | 0.1641 | 0.5887 | 0.3764 | 0.1083 | 3.4743 | 0.0005 | 0.004 | 0.3764 | 0.1641 | 0.5887 |
| | nsLearningYes | 0.5271 | 1.0959 | 0.8115 | 0.1451 | 5.5914 | 0 | 0 | -0.8115 | -1.0959 | -0.5271 |
| | socialYes | 0.2694 | 1.066 | 0.6677 | 0.2032 | 3.2867 | 0.001 | 0.008 | -0.6677 | -1.066 | -0.2694 |

#### 2.7 STATISTICS MAIN RESULTS: FEMALES

In the female dataset, we were unable to confirm our hypothesis in any of the domains investigated. In particular, in females with a history of ELA, we could not confirm changes in anxiety (HedgesG[95%CI] = .101[-.035,.236], z = 1.459, p=.59), memory after stressful learning (HedgesG[95%CI] = .192[.014, .37], z = 2.11, p=.14), memory after non-stressful learning (HedgesG[95%CI] = -.284[-.532,-.0355], z = 2.24, p=.1), or in social behavior (HedgesG[95%CI]=.011[-.405,.428], z = -.053, p=.957).

**Table legend**: ci.lb = lower boundary confidence interval; ci.ub = upper boundary confidence interval; effectsize = estimated Hedge's G; se = standard error of the estimated Hedge's G; z-value = z-value of the test; p-value = uncorrected p-value of the test; p-value_bonf = corrected p-value for family-wise comparison with Bonferroni; _Corr = refers to effectsize, ci.lb and ci.ub which have been flipped to ease interpretation (Method Section); sLearning = memory after stressful learning; nsLearning = memory after non-stessful learning; domainHit = statistics of experiments with multiple hits vs experiments without; domainNo = only experiments without multiple hits; domainYes = only experiments with multiple hits.

| | test | ci.lb | ci.ub | effectsize | se | Zvalue | Pvalue | Pvalue_bonf | effectsize_Corr | ci.lb_Corr | ci.ub_Corr |
|---|---|---|---|---|---|---|---|---|---|---|---|
| **Domains: main effects** | | | | | | | | | | | |
| | anxiety | -0.0347 | 0.2361 | 0.1007 | 0.0691 | 1.4587 | 0.1447 | 0.5788 | 0.1007 | -0.0347 | 0.2361 |
| | sLearning | 0.0138 | 0.3698 | 0.1918 | 0.0908 | 2.1123 | 0.0347 | 0.1388 | 0.1918 | 0.0138 | 0.3698 |
| | nsLearning | 0.0355 | 0.5321 | 0.2838 | 0.1267 | 2.2403 | 0.0251 | 0.1004 | -0.2838 | -0.5321 | -0.0355 |
| | social | -0.4276 | 0.405 | -0.0113 | 0.2124 | -0.0534 | 0.9574 | 1 | 0.0113 | -0.405 | 0.4276 |
| **Increased vulnerability: main effect** | | | | | | | | | | | |
| | hit | -0.0033 | 0.5965 | 0.2966 | 0.153 | 1.9387 | 0.0525 | 0.0525 | 0.2966 | -0.0033 | 0.5965 |
| **Effect increased vulnerability for each domain** | | | | | | | | | | | |
| | anxietyHit | -0.1384 | 0.3826 | 0.1221 | 0.1329 | 0.9188 | 0.3582 | 1 | 0.1221 | -0.1384 | 0.3826 |
| | sLearningHit | -0.001 | 0.6752 | 0.3371 | 0.1725 | 1.954 | 0.0507 | 0.2028 | 0.3371 | -0.001 | 0.6752 |
| | nsLearningHit | 0.0693 | 1.0611 | 0.5652 | 0.253 | 2.2338 | 0.0255 | 0.102 | -0.5652 | -1.0611 | -0.0693 |
| | socialHit | -0.6703 | 0.9941 | 0.1619 | 0.4246 | 0.3814 | 0.7029 | 1 | -0.1619 | -0.9941 | 0.6703 |
| **Posthocs** | | | | | | | | | | | |
| | anxietyNo | -0.1324 | 0.2118 | 0.0397 | 0.0878 | 0.4518 | 0.6514 | 1 | 0.0397 | -0.1324 | 0.2118 |
| | sLearningNo | -0.1816 | 0.228 | 0.0232 | 0.1045 | 0.2222 | 0.8242 | 1 | 0.0232 | -0.1816 | 0.228 |
| | nsLearningNo | -0.2997 | 0.3021 | 0.0012 | 0.1535 | 0.0076 | 0.9939 | 1 | -0.0012 | -0.3021 | 0.2997 |
| | socialNo | -0.5882 | 0.4036 | -0.0923 | 0.253 | -0.3649 | 0.7152 | 1 | 0.0923 | -0.4036 | 0.5882 |
| | anxietyYes | -0.0407 | 0.3643 | 0.1618 | 0.1033 | 1.5667 | 0.1172 | 0.9376 | 0.1618 | -0.0407 | 0.3643 |
| | sLearningYes | 0.0802 | 0.6404 | 0.3603 | 0.1429 | 2.5204 | 0.0117 | 0.0936 | 0.3603 | 0.0802 | 0.6404 |
| | nsLearningYes | 0.1719 | 0.9609 | 0.5664 | 0.2013 | 2.8133 | 0.0049 | 0.0392 | -0.5664 | -0.9609 | -0.1719 |
| | socialYes | -0.599 | 0.7382 | 0.0696 | 0.3411 | 0.2041 | 0.8383 | 1 | -0.0696 | -0.7382 | 0.599 |

#### 2.8 PUBLICATION BIAS RESULTS

Details on the tests used to evaluate the influence of publication bias are described in S1.8.

##### 2.8.1 MALES

Publication bias is evident from qualitative evaluation of funnel plot asymmetry (Figure A in S2.8), Egger's regression ($z = 7.501$, $p < .001$) and Begg's test ($z = 7.3961$, $p < .0001$). However, fail-safe file drawer analysis revealed that >7600000 unpublished, filed, or un-retrieved comparisons averaging null results would be required to bring the average unweighted effect size to non-significance. Similarly, trim-and-fill analysis (based on random effects meta-analysis) estimates that 0 studies are missing. These results suggest that although there is evidence of publication bias, the model seems not influenced by it as the interpretation of the results would not change.

##### 2.8.2 FEMALES

There is evidence of publication bias both from the funnel plot asymmetry (Figure B in S2.8), Egger's regression (based on random-effects meta-analysis, z = 2.329, p = 0.020) and Begg's test (z = 2.424, p = 0.015). However, fail-safe file drawer analysis was not performed as the overall meta-analysis was not significant (Q(8) = 14.384, p = .07). Trim-and-fill analysis (based on random effects meta-analysis) estimates that 0 studies are missing.

**Figure S2.8**. Funnel plots for publication bias evaluation in the A) males' and B) females' datasets.

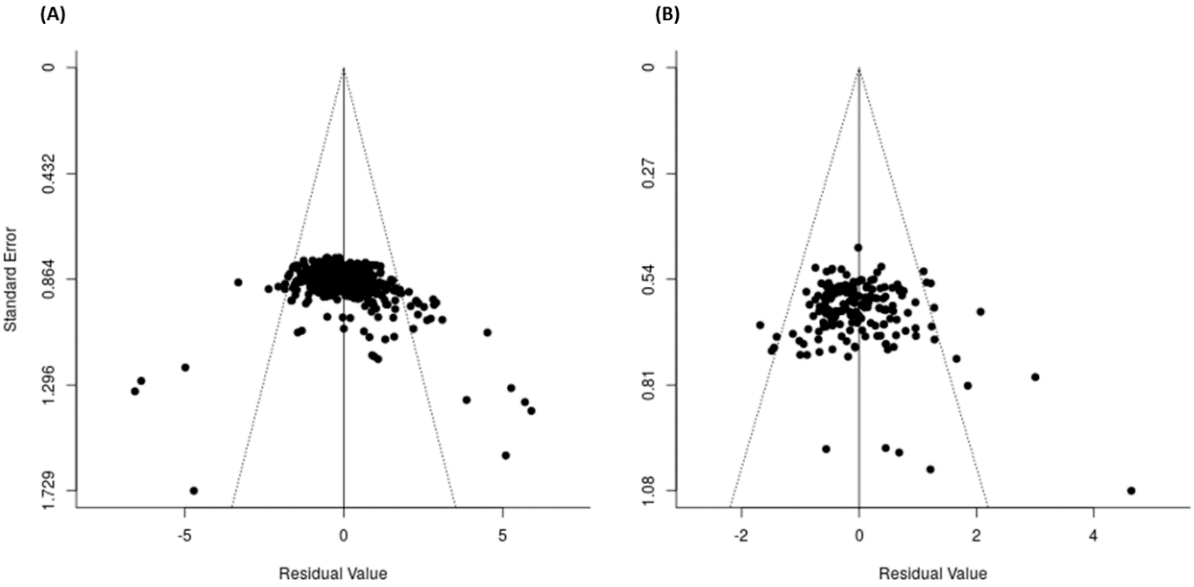

#### 2.9 SENSITIVITY ANALYSES

A summary of all sensitivity analyses performed can be found in supplementary material. Any researcher interested in analyzing sensitivity analyses in more detail is referred to the publicly available dataset and code used (https://osf.io/ra947/).

#### 2.10 STUDY OF DISTRIBUTION OF VARIANCE

In **males**, the within- ($\sigma_w^2$ = .296) and the between-variance component ($\sigma_b^2$ = .245) differed significantly from 0 (p<.000), which indicates that the variation in effect sizes is accounted for by differences within as well as between experiments. Conversely, in **females**, the between-experiment variance component ($\sigma_w^2$ = .000, p = 1.0) was negligible, while the within-variance component ($\sigma_b^2$ = .1838) differed significantly from 0 (p<.000). This indicates that the variation in effect sizes is accounted mainly by differences between experiments.

#### 2.11 DIRECTED EXPLORATION: METAFOREST PARTIAL DEPENDENCE PLOTS

Partial dependence plots are visualizations in which effect size is predicted as a function of the average over all other predictor variables. All partial dependence plots can be visualized by running the provided R-script (https://osf.io/ra947/).

Instructions:

1) Open the R project provided
2) Install any R package that might be missing.
3) Prepare environment by running the code section "Environment Preparation"
4) Prepare dataset for MetaForest analysis by running the code section "MetaForest: dataset preparation"
5) Perform MetaForest analysis by running the code section "MetaForest: tuning" (depending on your computer, it may take 30min-1h)
6) Save partial dependence plots in the Rproject folder by running the code section "MetaForest: plots".
